# SARS-CoV-2 Delta variant induces enhanced pathology and inflammatory responses in K18-hACE2 mice

**DOI:** 10.1101/2022.01.18.476863

**Authors:** Katherine S. Lee, Ting Y. Wong, Brynnan P. Russ, Alexander M. Horspool, Olivia A. Miller, Nathaniel A. Rader, Jerome P. Givi, Michael T. Winters, Zeriel YA. Wong, Holly A. Cyphert, James Denvir, Peter Stoilov, Mariette Barbier, Nadia R. Roan, Md. Shahrier Amin, Ivan Martinez, Justin R. Bevere, F. Heath Damron

## Abstract

The COVID-19 pandemic has been fueled by novel variants of concern (VOC) that have increased transmissibility, receptor binding affinity, and other properties that enhance disease. The goal of this study is to characterize unique pathogenesis of the Delta VOC strain in the K18-hACE2-mouse challenge model. Challenge studies suggested that the lethal dose of Delta was higher than Alpha or Beta strains. To characterize the differences in the Delta strain’s pathogenesis, a time-course experiment was performed to evaluate the overall host response to Alpha or Delta variant challenge. qRT-PCR analysis of Alpha- or Delta- challenged mice revealed no significant difference between viral RNA burden in the lung, nasal wash or brain. However, histopathological analysis revealed high lung tissue inflammation and cell infiltration following Delta- but not Alpha-challenge at day 6. Additionally, pro-inflammatory cytokines were highest at day 6 in Delta-challenged mice suggesting enhanced pneumonia. Total RNA-sequencing analysis of lungs comparing infected to uninfected mice revealed that Alpha-challenged mice have more total genes differentially activated, conversely, Delta-challenged mice have a higher magnitude of differential gene expression. Delta-challenged mice have increased interferon-dependent gene expression and IFN-γ production compared to Alpha. Analysis of TCR clonotypes suggested that Delta challenged mice have increased T-cell infiltration compared to Alpha challenged. Our data suggest that Delta has evolved to engage interferon responses in a manner that may enhance pathogenesis. The *in vivo* and *in silico* observations of this study underscore the need to conduct experiments with VOC strains to best model COVID-19 when evaluating therapeutics and vaccines.

**Importance:** The Delta variant of SARS-CoV-2 is known to be more transmissible and cause severe disease in human hosts due to mutations in its genome that are divergent from previous variants of concern (VOC). Our study evaluates the pathogenesis of Delta in the K18-hACE2 mouse model compared to the Alpha VOC. We observed that relative to Alpha, Delta challenge results in enhanced inflammation and tissue damage with stronger antiviral responses. These observations provide insight into Delta’s unique pathogenesis.

## Introduction

The COVID-19 pandemic is being perpetuated by the emergence of “Variants of Concern” (VOC), mutant strains of SARS-COV-2 with enhanced disease-causing abilities. Early in the pandemic, strains obtained the D614G mutation that enhanced binding of the spike protein to the host hACE2 receptor. The mutation quickly became dominant in the circulating strains around the world. Several potential variant strains were noted and monitored but few demonstrated any unique or enhanced differences in virulence or transmission. In September 2020, the B.1.1.7 strain was first identified in the United Kingdom and quickly spread to become the dominant variant worldwide in early 2021 (1). Later, in December 2020, two other variants, Beta (B.1.351) and Gamma (P.1) were identified in South Africa and Japan/Brazil, respectively (2–4). Both Beta and Gamma increased concerns because they decreased efficacy of several vaccines in clinical trials (4–10).

In addition to the effects on vaccine efficacy in clinical trials, there were also reports that Alpha and Beta variants were able to cause enhanced disease in humans (11–16) and animal studies with transgenic K18-hACE2-mice and hamsters supported those observations (17, 18). Before circulating VOC established their dominance, two doses of Pfizer-BioNTech’s mRNA vaccine, BNT162b2, exhibited 95% efficacy in protecting individuals from severe COVID-19 in retrospective cohort studies of COVID-19-associated hospitalizations in early 2021 (19). However, the efficacy of the BNT162b2 vaccine was slightly diminished in the context of the Alpha (93.7% efficacy) (20).

As Alpha, Beta, and Gamma continued to spread, the Delta variant (B.1.617.2) was identified in India, resulting in a massive surge of COVID-19 cases in the country (21). Unsurprisingly, Delta was able to cause breakthrough cases of infection within a portion of fully vaccinated people, decreasing the overall efficacy of ChAdOx1, mRNA-1273, or BNT162b2 vaccines (20,22–24). In March 2021, the Delta variant was detected in the United States, and reports of its increased transmissibility necessitated investigation of its threat to both unvaccinated and vaccinated populations. Key pathogenic mutations in the B.1.617 SARS-CoV-2 lineage, which includes the Delta (B.1.617.2) and Kappa (B.1.617.1) VOC, occur within the spike protein, which mediates viral attachment and entry into host cells via the ACE2 receptor (25, 26). These mutations have been reported to increase transmissibility within the population and decrease antibody neutralization (27). Delta does not harbor the N501Y substitution in the spike protein that was characteristic of the Alpha and Beta lineages but Delta does harbor the D614G substitution in spike, which contributes to the increased fitness and transmissibility of many VOC strains (28, 29). Some strains of Delta are reported to harbor the K417N mutation previously found in Beta and Gamma, which sparked debate over designating the strain as a new variant (30, 31). This mutant strain was labeled as “Delta Plus.” Additional mutations within the spike protein of Delta include P681R, which diverge from other VOC and help to enhance ACE2 receptor binding and cellular entry, contributing to reported increases in Delta’s pathogenicity (32).

To characterize the pathogenicity of Delta (utilizing strain B.1.617.2) and how it diverges from prior VOC (WA-1, Alpha, and Beta strains), we performed a time-course challenge study in K18-hACE2 transgenic mice, which were intranasally challenged with Alpha and Delta strains. Through analysis of viral load, tissue pathology, cytokine profiling, and total transcriptomics of the lung, we discovered that Delta causes increased interferon type I and II responses corresponding with greater lung inflammation when compared to Alpha. These data help to advance pre-clinical models of COVID-19 as well as serve as a benchmark to compare past and future VOC strains.

## Results

### The Delta Variant of SARS-CoV-2 has a greater LD100 than Alpha and Beta VOCs in K18-hACE2 mice

To date, most K18-hACE2 mouse or Golden Syrian hamster studies have utilized ancestral viral strains of SARS-CoV-2 (e.g., WA-1, D614G, B.1 Wuhan), and few studies have been performed with emergent VOC (2,17,18,33–39). Early studies in K18-hACE-mice with a variety of viral strains used a wide range of challenge doses from as low as 100 PFU to as high as 10^5^ PFU (40–42). In pilot studies with the Alpha and Beta variants, we evaluated 10^5^ PFU (high dose) and observed mortality as early as day 4 post challenge, suggesting that VOCs have enhanced virulence in K18-hACE-mice compared to WA-1 ancestral strain (data not shown). We then aimed to identify an appropriate challenge dose for WA-1, Alpha, Beta, and Delta VOC that could cause symptomatic disease for appropriate comparisons between the subtle aspects of variant pathogenicity masked by lethal challenge doses. At the low dose of 10^3^ PFU, WA-1 challenge only resulted in morbidity of 50% of the mice (Fig. 1A). We observed that a challenge dose of 10^3^ PFU using Alpha or Beta resulted in 80% and 100% mortality, respectively by day 7 post challenge. Although this comparison between Alpha and Beta challenge at 10^3^ PFU is not statistically significant (*P=* 0.6452), there is a significant difference between the survival of WA-1 challenged mice compared to Beta at the 10^3^ PFU dose, where WA-1 elicits higher survival (*P=* 0.0446). Despite the observations of enhanced virulence of Delta in humans, 10^3^ PFU challenge of K18-hACE2-mice mirrored WA-1 survival more so than Alpha or Beta (Fig. 1A). As expected, increasing the challenge dose to 10^4^ PFU decreased survival of K18-hACE2-mice for all strains (Fig. 1B). At the 10^4^ PFU challenge dose, Delta resulted in 0% survival (Delta vs WA-1 *P*= 0.0002) (Fig. 1A-B). The 10^4^ PFU challenge dose also shortened the time to morbidity for Beta as well as Alpha (WA-1 vs Alpha *P*=0.0008; WA-1 vs Beta *P*=0.0122). As we suspected from an ancestral strain, total morbidity of WA-1 infected groups was not observed at the either dose. Using a disease scoring system established previously (T.Y.Wong, K. S. Lee, et al., JVI accepted for publication) we observed higher average disease scores for Alpha and Beta than Delta in K18-hACE2 mice intranasally challenged with 10^3^ PFU (Alpha vs Delta *P*=0.0355; Beta vs Delta *P*=0.0039) (Fig. 1C-D). Interestingly, Delta’s observable disease phenotypes remained low by day 6 when Alpha- and Beta- challenged mice started to develop greater observable disease phenotypes consistent with morbidity (Alpha vs Delta *P*=0.0056). Delta’s disease phenotypes were scored higher following 10^4^ PFU challenge, where they progressed in a pattern similar to Alpha. Delta-challenged mice received higher average disease scores than Beta by day 6 post challenge (Fig. 1D). These experiments suggested that Delta compared to WA-1, Alpha, and Beta We reasoned that these differences warranted further investigation.

**Figure 1.**
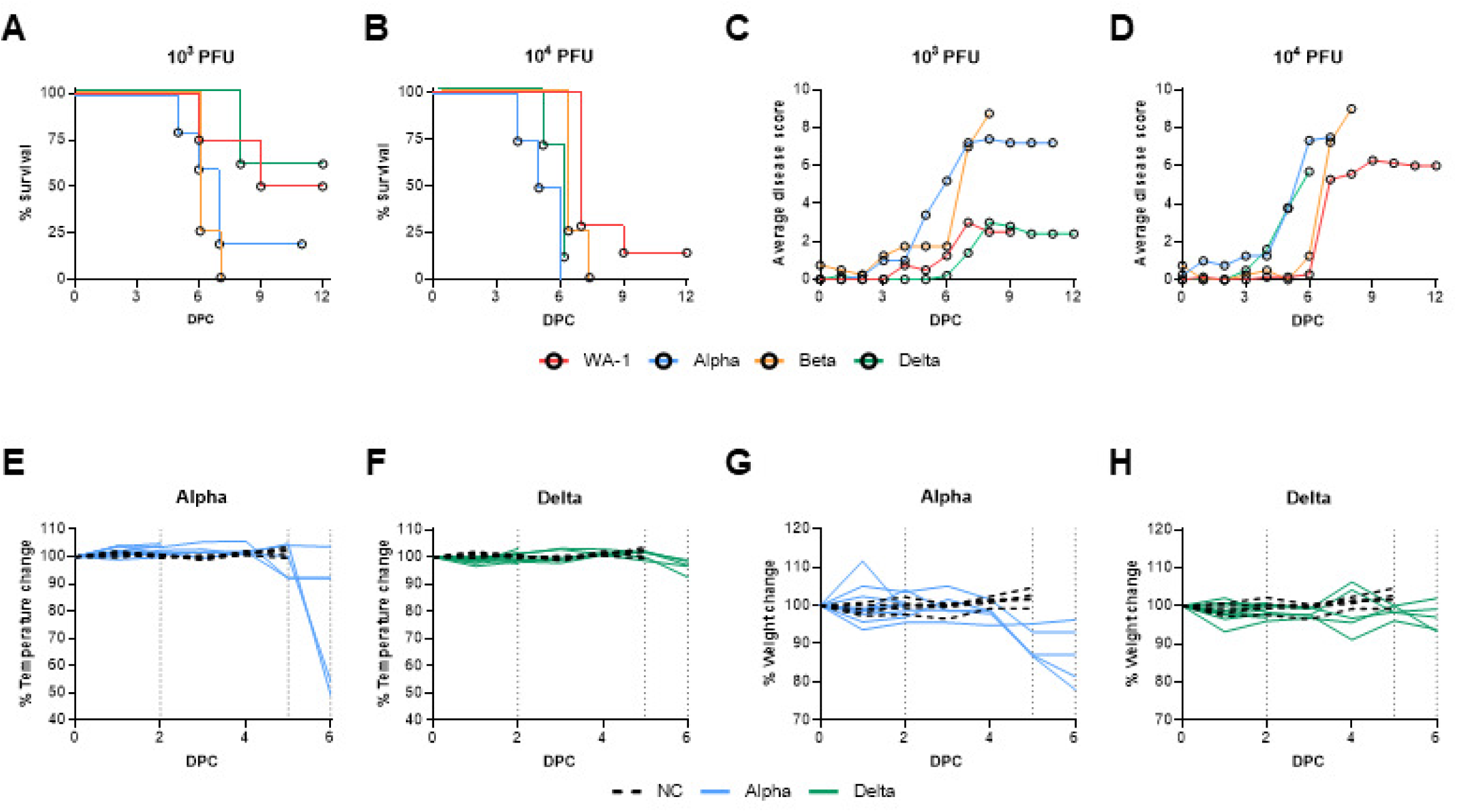
Survival, disease scores, weight, and temperature of K19-hACE2-mice challenged with WA-1, Alpha, Beta, and Delta strains. K18-hACE2 transgenic mice were intranasally challenged with 1×10^3^ or 1×10^4^ PFU WA-1, Alpha (B.1.1.7), Beta (B.1.351) or Delta (B.1.617) SARS-CoV-2, or mock-challenged with PBS (NC). Post-challenge, mouse survival (A, B) (n=4 for 1×10^3^ WA-1, n=7 for 1×10^4^ WA-1, n= 3 for 1×10^3^ Alpha, n=4 for 1×10^4^ Alpha, n=4 for 1×10^3^ Beta, n=4 for 1×10^4^ Beta, n=5 for 1×10^3^ Delta and n=10 for 1×10^4^ Delta) and disease scores (C, D) were evaluated. Based on experimental results, mice challenged with Alpha (n=10 split between timepoints) and Delta (n=10 split between timepoints) or mock-challenged with PBS (n=5), were evaluated at day 2 and day 6 post-challenge to measure changes in bodyweight (E, F) and body rectal temperature (G, H). Dotted lines indicate euthanasia points on day 2, day 5 (euthanasia of mock-challenge controls) and day 6. NC=no challenge.

### Mice challenged with of Alpha or Delta experience unique disease manifestation

To gain additional insights into the differences in pathogenesis of Alpha and Delta in K18 mice, we focused on a challenge dose of 10^3^ PFU which allows for differential disease manifestation between Alpha and Delta. Rectal temperature and weight loss were monitored in mice challenged with 10^3^ PFU of Alpha or Delta. In agreement with their disease scores, Alpha challenged mice had significant loss of body temperature (hypothermia) and weight loss between days 5 and 6 post challenge (Fig. 1E-F). However, Delta-challenged mice maintained body temperature and did not experience weight loss (Fig. 1G-H). At day 6 post challenge, Alpha challenged mice had reached morbidity based on disease scoring and required euthanasia as a humane endpoint. However, at day 6 Delta challenged mice remained below euthanasia criteria. Collectively, these data demonstrate that Alpha and Delta cause distinct disease profiles in K18-hACE2-mice.

### Mice challenged with Delta have similar levels of viral RNA burden in the nares, and lungs but lower amounts of viral RNA in the brain

It has been previously reported that SARS-CoV-2 viral burden is highest two days after challenge (43) and based on our data we recognize that Alpha variant will cause morbidity at day 6 post-challenge; however, Delta variant challenge will not cause the same disease, or survival. Therefore, we aimed to evaluate viral RNA burden at day 2 and 6 post challenge for Alpha or Delta challenge mice. At euthanasia on day 2 and 6 the lung, brain, and nasal wash fluid were collected from challenged mice to quantify viral RNA burden via nucleocapsid qRT-PCR. We observed that viral RNA burden was remarkably similar between Alpha and Delta in the lung at both time points (Fig. 2A). In the nasal wash, viral load was also detectable at similar levels for both Alpha and Delta at day 2. In contrast to the lung, the viral RNA burden decreased to undetectable levels at day 6 for both VOC (Fig. 2B). Surprisingly, challenge with Delta did not lead to detectable levels of viral RNA in the brain at either day 2 or day 6. In contrast, mice infected with Alpha exhibited low RNA levels at day 2 and a 2-fold increase at day 6, suggesting viral replication in the brain over time (Fig. 2C). These observations suggest that Delta replicates efficiently in the airway in a manner similar to Alpha but Delta appears to lack the brain localization previously observed in K18-hACE2 mice (43–46).

**Figure 2.**
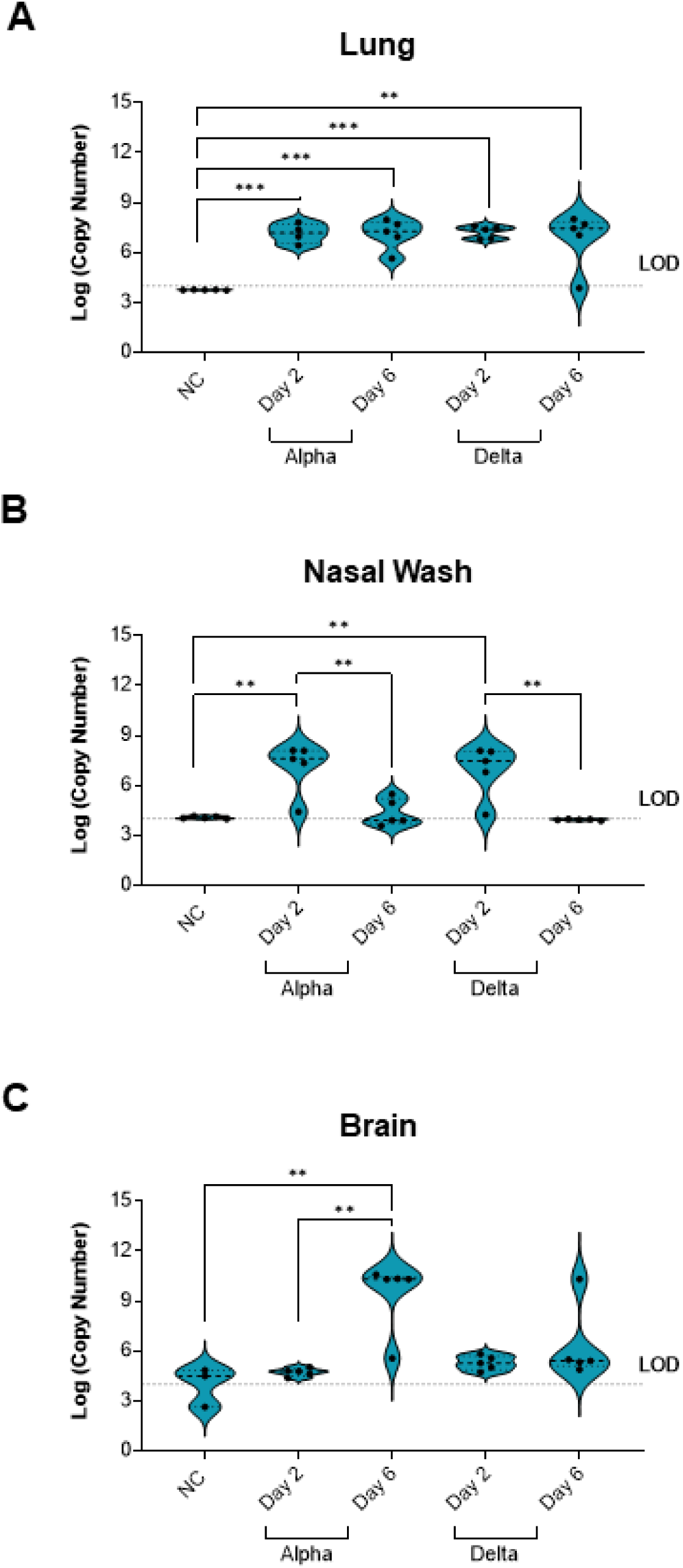
Viral RNA burden of disease-associated tissues was quantified using nucleocapsid qPCR. Viral RNA was detectable in the lung tissue of challenged mice from both variants at both timepoints compared to no challenge (One-Way ANOVA, *P=*0.0008; 0.0004; 0.0003; 0.0013) (A). In Nasal Wash, viral RNA burden increased from both variants at Day 2 compared to no challenge (One-Way ANOVA, *P=*0.0016 Alpha; *P=*0.0030 Delta) and was decreased at Day 6 for both variants from copy numbers at Day 2 (One-Way ANOVA, *P*=0.0043 Alpha; *P=*0.0019 Delta) (B). Alpha challenged mice have increased detectable viral RNA in the brain at Day 6 compared to Day 2 (One-Way ANOVA, *P=*0.0012) while Delta challenge results in no viral RNA burden in the brain (C). Dashed lines indicate limit of detection via qPCR. NC=no challenge.

### Mice challenged with Delta experience significant inflammation in the lung tissue

Based on qRT-PCR data showing viral RNA burden in the lung but not brain of 10^3^ PFU Delta-challenged mice, we hypothesized that relative to other Alpha, Delta may cause robust pneumonia and less encephalitis in K18-hACE2-mice. To test this, we performed histopathological analysis on the lungs collected at days 2 and 6 to characterize the pneumonia caused by Alpha or Delta challenge. At two days following challenge, inflammation, and recruitment of inflammatory cells could already be observed in the Delta- but not Alpha-challenged mice when compared to the lungs of uninfected mice (Fig. 3A-B). At this time, a pathologist identified that the inflammatory infiltrate was composed predominantly of lymphocytes and occasional histiocytes. By day 6, an increase in marked perivascular inflammation and margination was measured in Delta-(36.86 marginating lymphocytes per mm length of endothelium) and Alpha-challenged mice (12.19 marginating lymphocytes per mm length of endothelium) (unpaired t-test *P=*0.026) (Fig 3C-D). In Alpha lungs, less total vessels were identifiable within areas of inflammation, awarding these samples a margination score of zero cells/mm of vessel. Overall, less inflammation occurred in Alpha lungs. At day 6, only 1% of the lung area featured inflammation in Alpha, compared to 20% for Delta-challenged mice (unpaired t-test *P=*0.003) (Fig 3E). Within these tissue areas (the airway, alveoli, and thin mucous layer), the presence of Delta virions was identified using electron microscopy (Fig 3F-G). Collectively, these data suggest Delta challenged mice have increased cellular responses and inflammation compared to Alpha challenged K18-hACE-mice.

**Figure 3.**
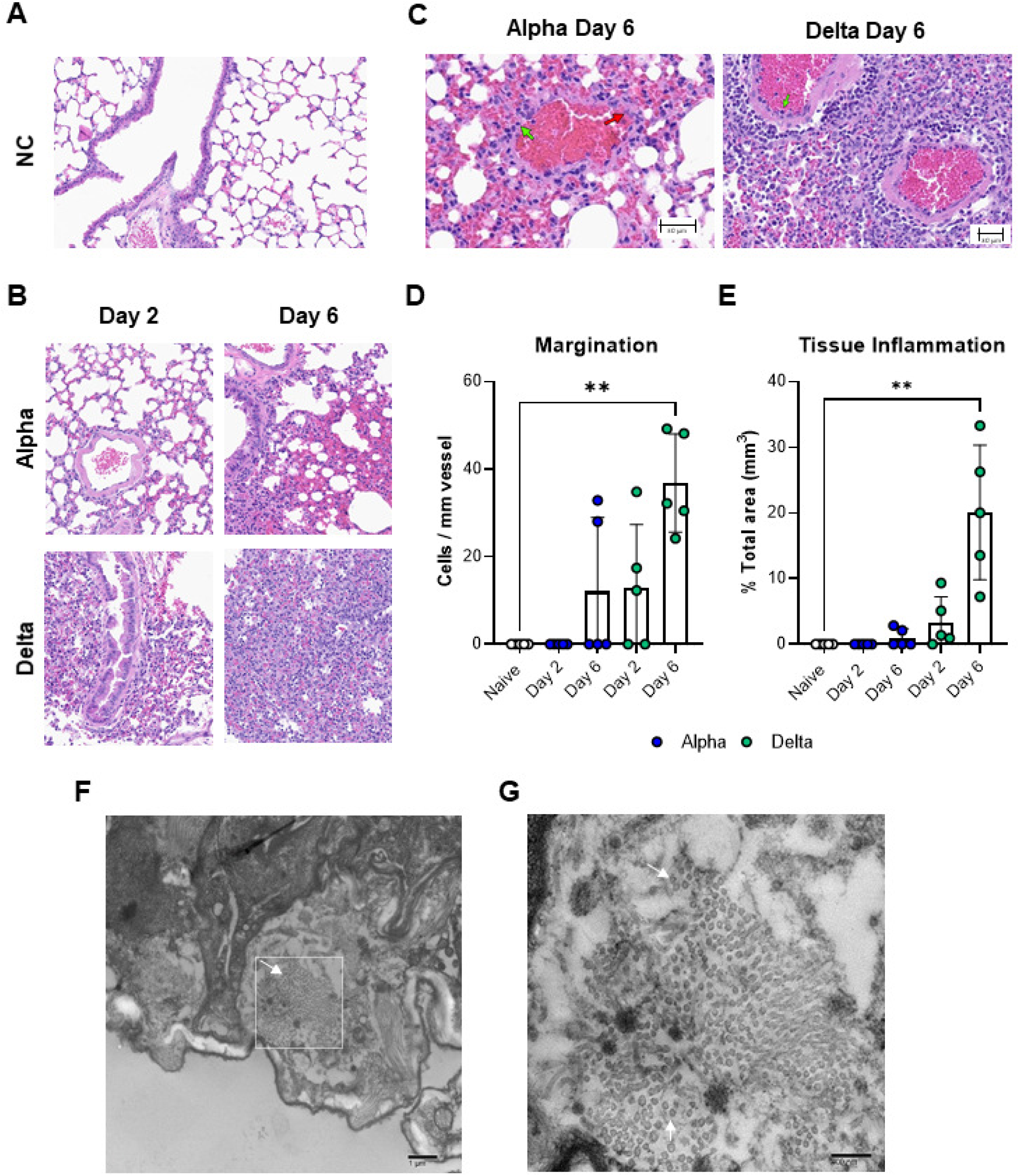
Histopathological and electron microscopy analysis of Alpha or Delta challenged K18-hACE2-mouse lungs. 20X images demonstrating inflammation in the lung tissue of Alpha and Delta SARS-CoV-2 challenged K18-hACE2 at Day 2 or 6 post-challenge (B) compared to no challenge (A). 20X images reveal margination at blood vessels within areas of inflammation (green arrows indicate counted marginating cells, red arrows indicate tissue-resident non-marginating cells)(C). Marginating inflammatory cells were counted within the tissue and were found to be increased in Delta challenge at Day 6 (Kruskal Wallis One-Way ANOVA, *P=*0.0097) (D). Regional tissue inflammation was quantified as % total area of analyzed tissue (mm^3^) and was highest in Delta at Day 6 compared to no challenge (Kruskal Wallis One-Way ANOVA, *P*=0.0012) (E). Error bars represent SD. Electron microscopy images displayed distribution of Delta virus particles in 1×10^3^ PFU challenged mouse lung tissue (F,G). White arrows identify virions.

### Pro-inflammatory cytokines are increased in mice challenged with Delta at day 6

Histopathological analysis revealed significant inflammation in the lungs of Delta-challenged mice at day 6 compared to Alpha-challenged mice despite equal viral RNA burden. To better understand differences in the inflammatory profile of these tissues, pro-inflammatory cytokine levels in lung supernatant at day 2 or 6 were quantified to profile strain specific variations in the “cytokine storm” (47). Insignificant amounts of cytokine were produced in the lungs of mice challenged with Alpha or Delta at day 2 (Fig. 4). Alpha-challenged mice produced low amounts of IL-1β and CXCL13 compared to uninfected mice at day 2 or 6. By contrast, the cytokine response to Delta challenge at day 6 compared to uninfected revealed high levels of IL-1β, TNFα, and CXCL10 (*P=*0.0462*, P=*0.0268*, P=*0.009) (Fig. 4). The inflammatory response within the lungs of mice challenged with Delta was elevated compared to Alpha. To gain insight into the specific mechanisms of the immunological host response, we performed RNAseq to characterize the total transcriptional profile of Alpha and Delta challenged K18-hACE2 mouse lungs.

**Figure 4.**
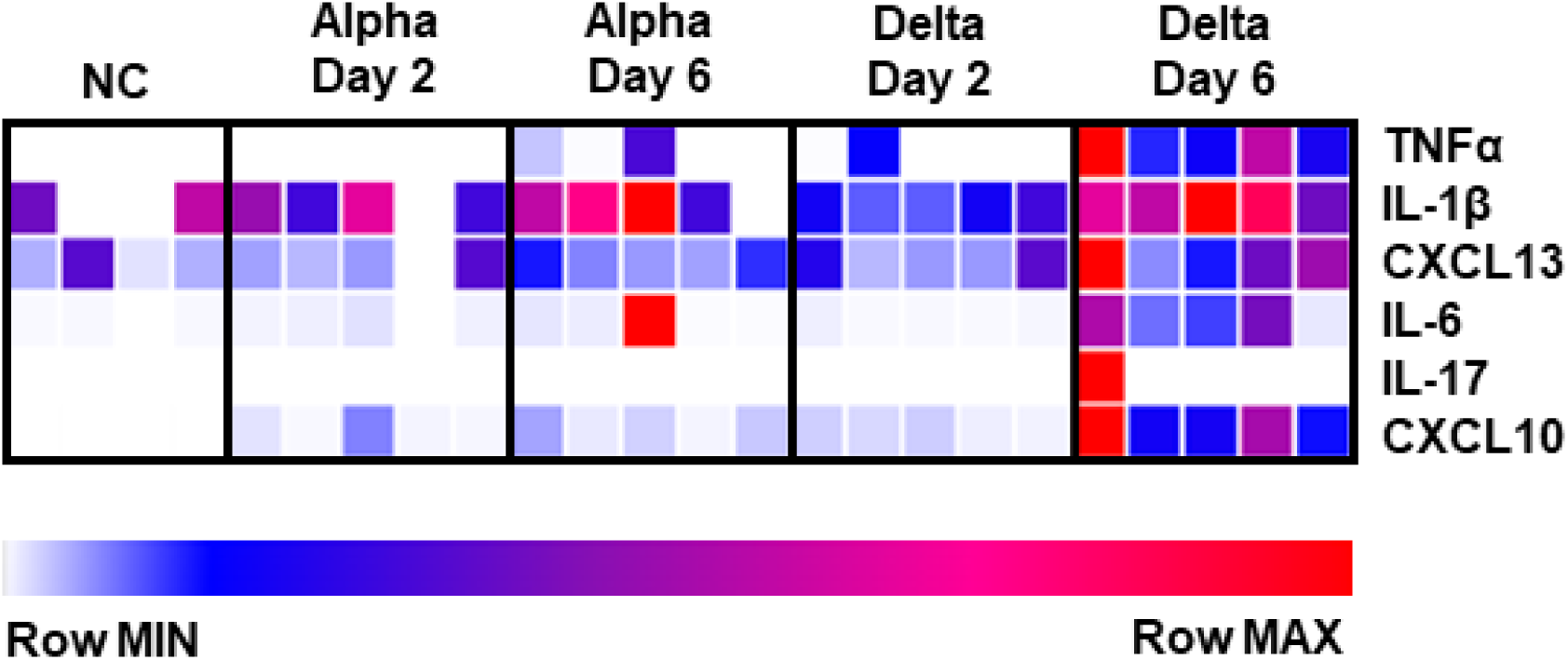
Analysis of cytokines production in lung supernatants from Alpha or Delta challenged K18-hACE2 transgenic mice at 2 or 6 days post-challenge. Non-challenged, Alpha or Delta challenged lung supernatants were used to determine local cytokine production in response to challenge. Concentrations (pg/mL) were graphed using Morpheus to reveal relative levels of cytokines compared to non-challenged lungs.

### RNAseq analysis of lung tissues from mice challenged with Alpha or Delta shows no difference in viral sgRNA

We utilized RNA that was isolated from the lung tissue of challenged and uninfected mice at 2- and 6-days post-challenge for Illumina transcriptomic analysis. RNA sequencing reads were first mapped to the SARS-CoV-2 viral genome to quantify the expression levels of viral genes over the course of infection (Fig. 5A). This revealed no significant difference between the two strains (Fig. 5B-C) which directly supported the qRT-PCR analysis (Fig. 2). It is known that SARS-CoV-2 infection results in down-regulation of hACE2 expression(48, 49). Therefore, we mapped RNA reads to the hACE gene to analysis this effect. Although not significant, a trend of lowered hACE2 expression occurred in Alpha challenged mice, whereas Delta challenged mice appeared to experience only slightly reduced hACE2 expression (Fig. 5D). These observations together with qRT-PCR nucleocapsid data described above, suggest that with equal viral burden (same challenge dose and equal viral RNA burden), the differences in LD100 of Alpha and Delta as well as the disease phenotypes are likely related to differences in host response to VOC and not overall viral load.

**Figure 5.**
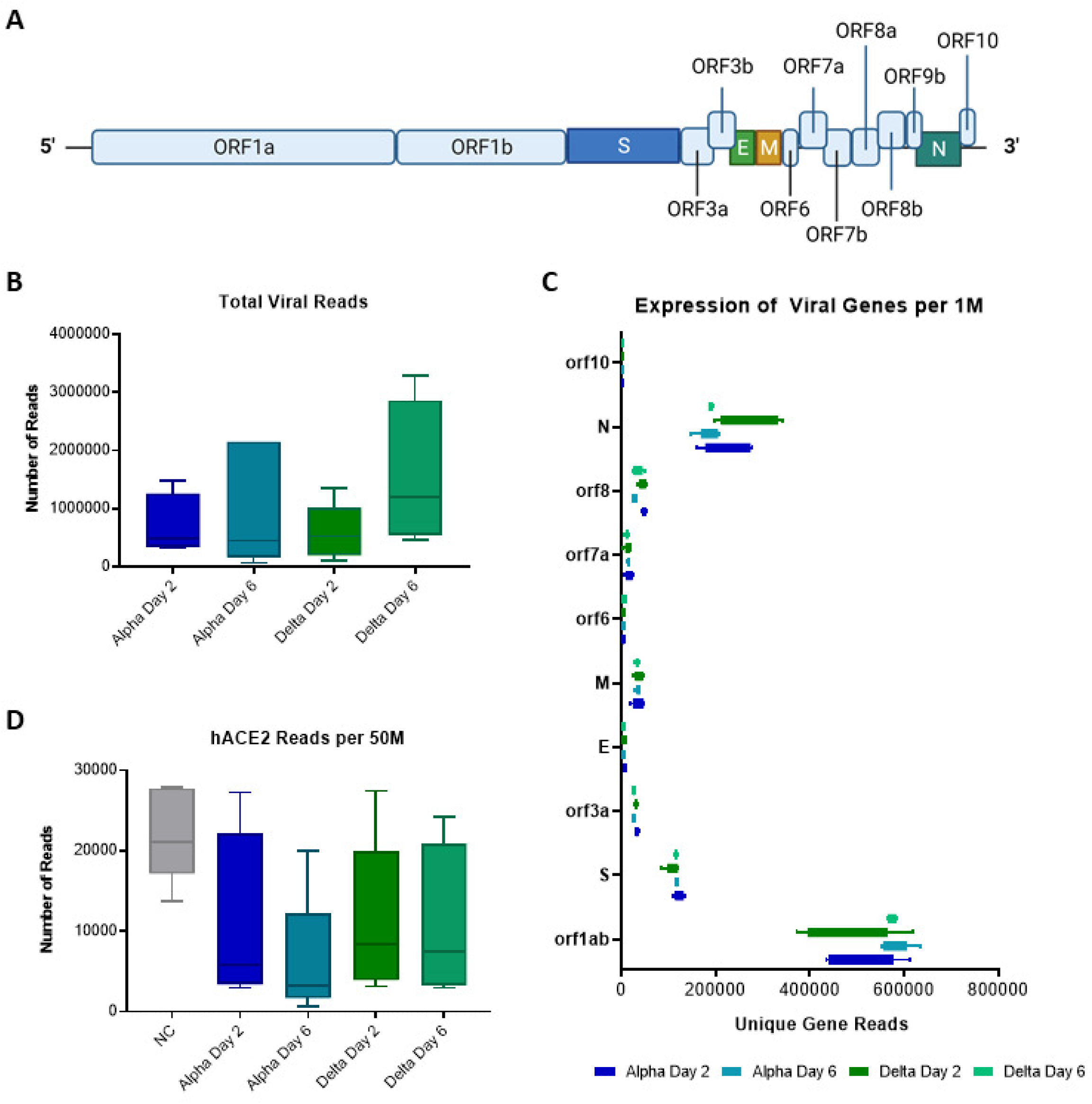
RNAseq analysis of SARS-CoV-2 genes in challenged mouse lungs. Schematic of the SARS-CoV-2 genome (A). Total viral read counts in each group (B) as well as quantification of viral reads per 1M RNAseq reads obtained (C). Reads of human ACE2 gene of the K18-hACE2-mice were quantified per 50M reads obtained per lung sample (D).

### Delta challenge causes a higher magnitude of host gene expression change compared to Alpha

To compare the host responses of Delta- and Alpha-challenged mice, we mapped lung RNA sequencing reads to the mouse genome. We observed that Alpha or Delta challenge in K18-hACE2 mice result in distinct transcriptional changes. Two days after challenge with Alpha, 2,413 statistically significant genes (FDR *P* ≤ 0.04) were differentially expressed compared to uninfected mice, a difference which increases more than two-fold to 5,664 genes at day 6. In Delta groups, differential expression encompasses 3,048 and 4,489 genes respectively. Venn diagrams of the gene sets comparing by variant and time-points illustrate overlaps in gene sets that are either activated or repressed. Both activated and repressed gene sets increase from day 2 to 6 (Fig. 6A). Further filtration of these sets of genes to identify those that are uniquely affected by VOC challenge and timepoint revealed that Alpha challenge results in more genes differentially regulated at day 6 than Delta challenge (Fig. 6B). A set of core-activated genes was identified through a four-way comparison of VOC and timepoint. Alpha and Delta challenge both activate a similar set of host genes despite their different disease pathology. After identifying the commonalities in our data set, we sought to identify how host transcriptomics differ between the groups. When the fold changes of this gene set were graphed on a curve according to their respective values at 6 days post-Delta challenge, it became clear that expression of these core genes follows a similar trend (Fig. 6C).

**Figure 6.**
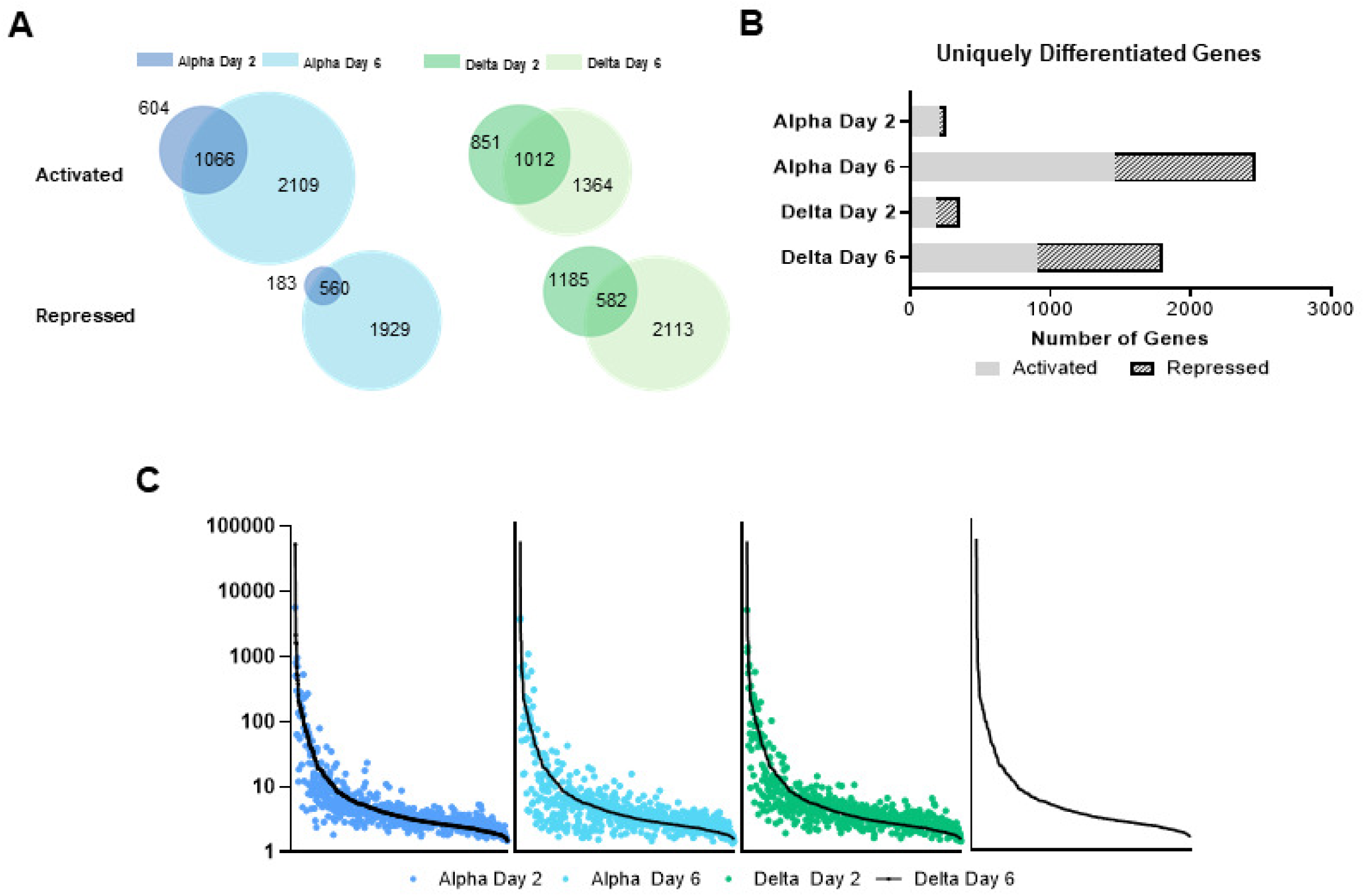
Total transcriptomic analysis of mouse genes in non-challenged mice compared to Alpha and Delta challenged mice at days 2 and 6. RNAseq reads from challenged mice were compared to non-challenged mice to determine differential gene expression (A). Genes that were statistically significant are indicated in relation to challenge day and their relative expression as being activated or repressed. Venn diagrams are shown to illustrate the amount of overlap in significant genes between days 2 and 6. The total number of unique differentially expressed genes are represented per each challenge day and relative expression (B). The highest differentially activated genes are show per each group with fold change relative to non-challenged control RNAseq reads (C). Gene expression of activated genes are ranked by fold change in Delta day 6 mice and then shown in relation to both strains and days post challenge (D).

### GO analysis revealed unique systems of enriched genes in mice challenged with Delta

Gene Ontology (GO) term analysis was performed to systematically group transcriptomic observations to identify specific pathways and gene systems whose expression changes in response to Alpha and Delta challenge. Relative to Alpha-challenged mice, Delta-challenged mice had a higher number of unique GO terms that were enriched at day 6 (Fig. 7A). To categorize the specifically enriched terms in Delta at day 6, pathway enrichment ratios were calculated for the top 30 GO terms (Fig. 7B). The expression of genes that were annotated with GO terms pertaining to immune responses, anti-viral, or lymphocyte recruitment were increased, as were genes of pathways associated with T cells and responses to IFN-β (Fig. 7B). Due to the fact that GO term analysis identified T cell response genes to be enriched in Delta challenged mice, we performed T cell receptor (TCR) clonotype analysis using RNAseq reads. Within the uninfected mice, we detected ∼50 total clonotypes. In Alpha-challenged mice, the number of clones were decreased relative to uninfected at both day 2 and 6. By contrast, Delta-challenged mice had decreased clones at day 2, but we observed a 6-fold increase in TCR clones at day 6 (Fig. 7C). In counts of unique TCR clonotypes, the same trends were observed: at day 6, Delta challenge had the most total unique TCR clones compared to non-challenged mice, day 6 post-Alpha challenged mice, and day 2 post Delta challenged mice (Fig. 7D). These data suggest that Delta may induce more robust T cell response.

**Figure 7.**
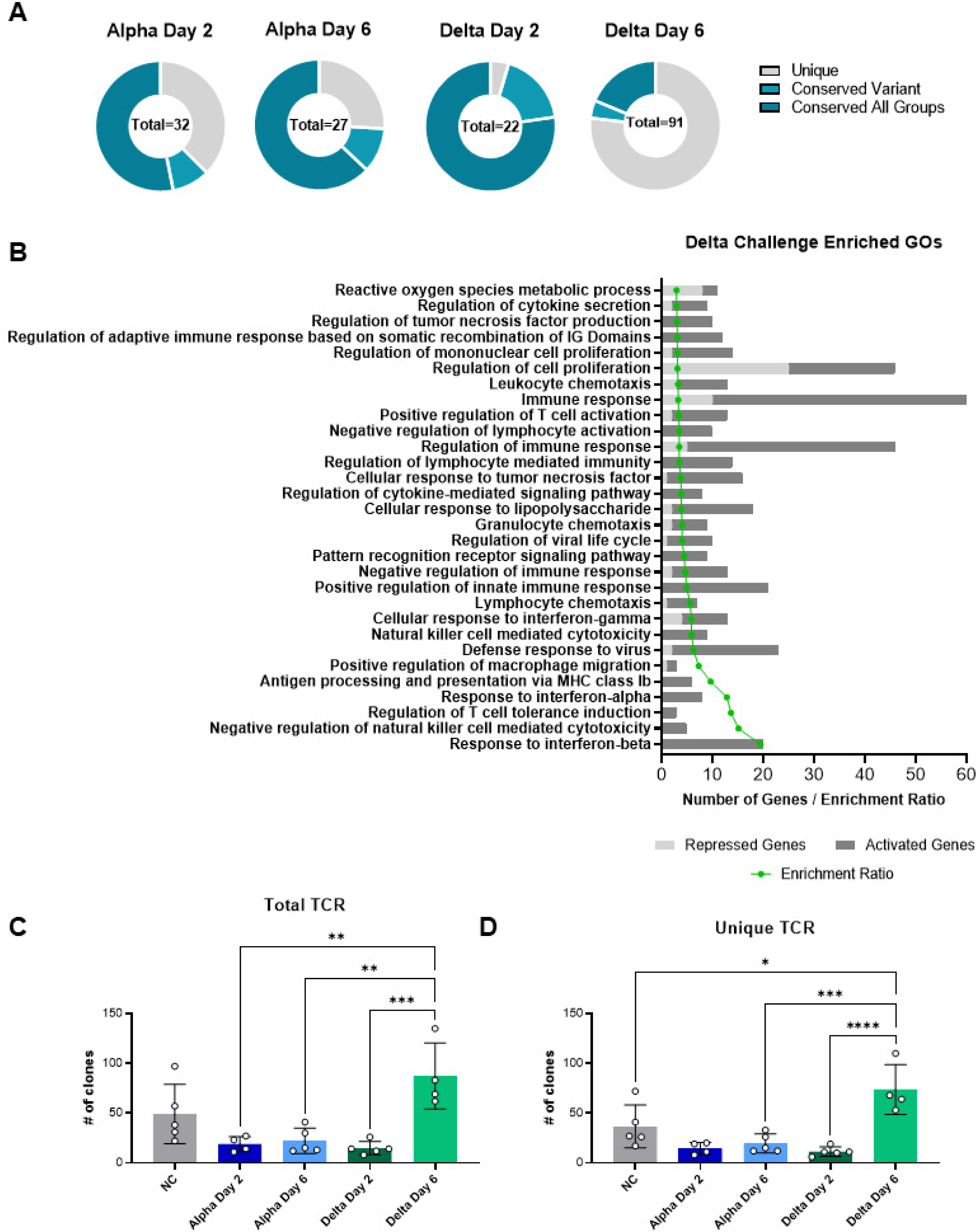
Gene ontology analysis of systems of genes in challenged mice. Circle plots indicate the number of unique or conserved sets of genes that were found to be enriched per each group (A). The top 30 enriched GO terms are shown and plotted per their relative level of expression and enrichment ratio (B). TCR clonotypes were mapped and represented as Total clones (C) or unique clones per unique CDR3 sequence (D). Total TCR clones (*P*=0.0005) and unique clones were increased between Delta day 2 and day 6 (*P*<0.0001).

### Delta challenge causes increases interferon dependent gene expression

Gene expression analysis focused by GO terms revealed Delta challenge induced expression of interferon response genes (Fig. 7). Interferon is an import regulator of the antiviral response in humans. It’s variable expression in response to specific variants exemplifies the differences in the host response. TGTP1 and IFI211, two interferon related genes, are the highest-expressed genes in Delta at day 6 at a fold change of 52,725 or 11,733 higher than in uninfected mice. In Alpha at day 6 these genes are 3,367 and 591-fold higher than uninfected. Due to the important role of interferons in the anti-viral response to SARS-CoV-2 which may be differentially engaged by VOC, we continued to compare the top 20 genes involved in general interferon responses. Total gene counts in lung RNA from each experimental group are shown in Fig. 8A. Delta day 6 shows high interferon dependent gene expression as well as the highest fold change difference (Fig. 8B). Collectively, these data suggest Delta challenge results in higher interferon response. To test that hypothesis, we directly measured type I and II interferons in lung supernatants and serum. Modest IFN-α or IFN-β were observed in both lung and serum (IFN-α: *P*=0.0012 Delta day 6 vs No Challenge in lung supernatant; IFN-β: *P*=0.0474 Alpha Day 2 vs No Challenge in serum); however, Delta challenge resulted in high IFN-γ at day 6 (*P*=0.0265 in lung supernatant, *P*=0.0103 in serum compared to no challenge) (Fig. 8C-D).

**Figure 8.**
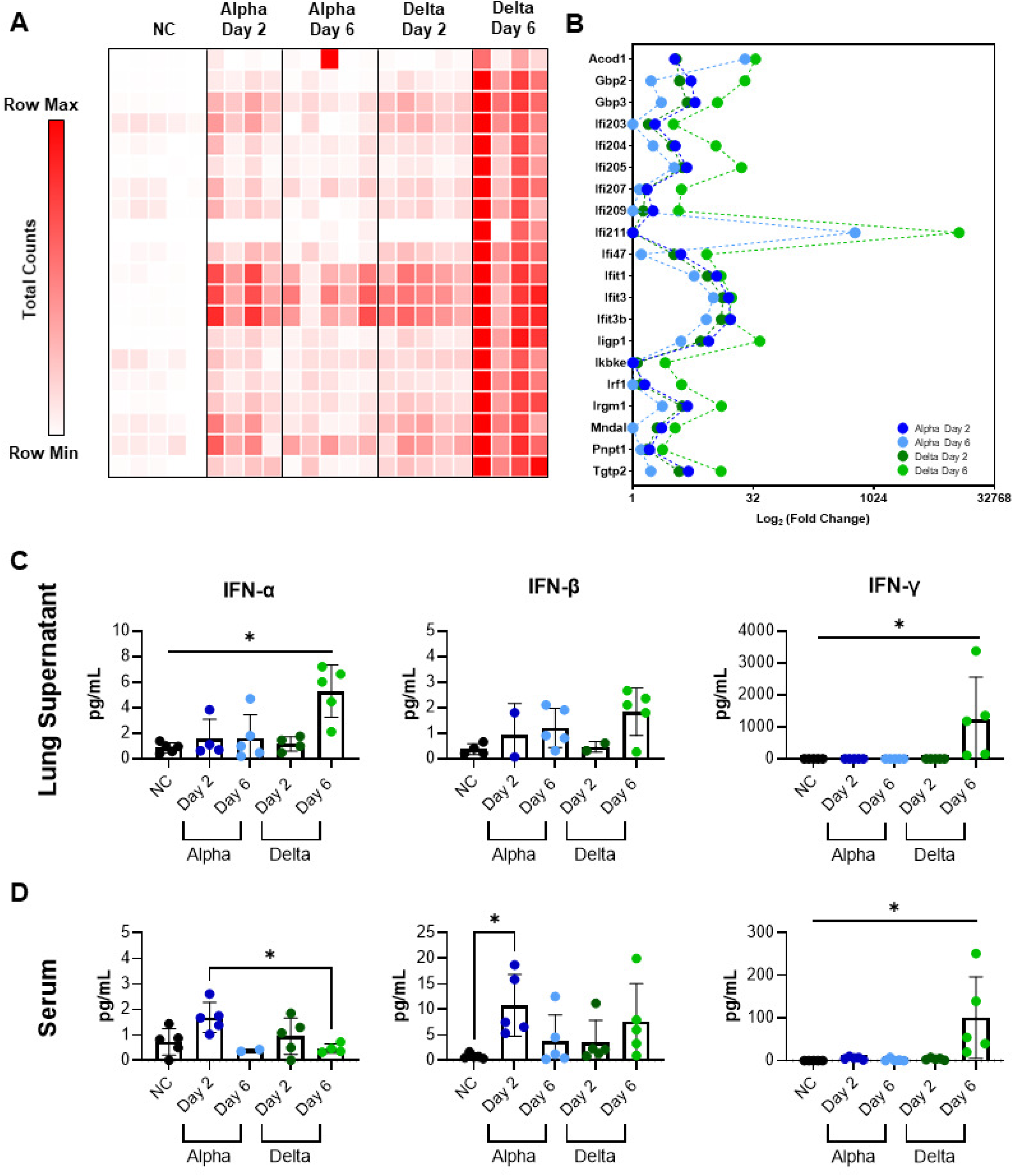
Analysis of interferon dependent gene expression and production of type I and type II interferon in lungs and serum of K18-hACE2-mice. Total RNA counts per interferon dependent gene are shown per each mouse, time-point, and variant challenge group (A). Each square represents the total RNA counts per gene from one mouse. Interferon dependent genes are shown with relative fold change plotted (B). Interferon alpha, beta, and gamma production in lung supernatant (C) and serum (D) of K18-hACE2-mice.

## Discussion

In this study, we modeled the pathogenesis of SARS-CoV-2 variants of concern in the K18-hACE2 mouse challenge model. Based on our observations that the Delta variant differs in disease manifestation from Alpha and Beta VOC-challenge, we performed an experiment to compare Alpha- and Delta-challenged mice at disease-significant timepoints. Despite a similar viral RNA burden in the respiratory tract at day 2 and 6, Delta causes more significant inflammation in the lungs than Alpha as infection progresses. The host response also varies over time, with Delta causing increased antiviral cytokine production, specifically IFN-γ, at day 6. GO term analysis of Delta-challenged mice suggested a greater number of immunological pathways that are implicated in the variant’s pathogenesis compared to Alpha, further indicating the unique nature of host response to individual variants.

Severe COVID-19 is perpetuated by dysregulated production of pro-inflammatory cytokines, including type I and II interferons, in a “cytokine storm” pathology (47,50,51). Interferon-stimulated genes (ISG) in patient lung tissues implicate the role of interferon in SARS-CoV-2 pathogenesis, but it has not been determined if the cytokines contribute more to suppressing viral replication, or suppressing the immune response through their anti-inflammatory abilities, contributing to greater disease (50, 51). Therefore, it is important to investigate how VOC differentially engage interferon production and subsequently the expression of interferon stimulated genes, during infection. Delta challenge causes a significant increase in lung and serum IFNγ which is greater than what has been previously reported in K18-hACE2 mice using different challenge strains such as ancestral WA-1 (30,31,32). IFNγ is a potent antiviral cytokine that occurs later in the timeline of viral challenge and may not be observable in widely used COVID-19 mouse models because of short survival lengths (52). IFNγ production is typically associated with the activation of Natural Killer cells, innate lymphoid cells, or Th1-like adaptive immune cells (44). Its high concentrations in the lung of Delta mice could corroborate greater recruitment of these cell populations identified in histopathology data as infiltrating cells (Fig. 3). Together the observations of this study suggest that Delta challenge causes enhanced innate immune responses compared to Alpha.

Surprisingly we observed that the viral RNA burden of the respiratory tract was identical per day per variant in the K18-hACE2-mice, as determined by both RT-qPCR and RNAseq, a clear question remains: why does Delta induce more interferon responses? One possible option that could be explored is the interaction of viral sgRNAs in regulating host gene expression. Open Reading Frames (ORFs) of SARS-CoV-2 have been implicated in controlling the host anti-viral response (53). Studies have reported that the increase in viral burden due to Alpha leads to increases in some but not all sub-genomic RNAs from the genomic regions harboring variant-characteristic mutations (54). Another study links the expression of sub-genomic RNAs in Alpha infection, specifically ORF9b, to antagonism of the host immune response and suppression of ISGs which would allow for more efficient viral replication and enhanced transmission (55). Although viral gene reads in our analysis show no significant differences between Alpha and Delta sgRNAs, it is possible that our selected timepoints were too late to detect the differences previously reported at 10 and 24 hours post challenge, or that our challenge dose was too low. Still, our abundant transcriptomic data suggest an extensive list of gene pathways that were enriched during Delta but not Alpha infection.

We observed that 10^3^ PFU challenge with Delta variant does not cause the dramatic weight loss and hypothermia reported in models of Alpha and Beta challenge (17). One well known caveat of the K18-hACE2-mouse model is the fact that WA-1, Alpha and Beta strains enter the brain (43,45,46,56–62). However, it appears that Delta variant remains more localized in the lungs of challenged mice (Fig. 3) (63). It’s generally accepted that the K18-hACE2-mice reach morbidity due to brain infection, supported by manifestations of disease including hypothermia, and inflammatory profiles. It is possible that the absence of brain infection due to Delta allows for an increased survival time, leaving more time for the development of severe infection and pneumonia in the lung, which could make Delta a better strain for modeling the respiratory phenotypes of COVID-19 disease in K18-hACE2 mice.

We appreciate that the K18-hACE2-mouse model has caveats to consider. The transgenic model expressing human ACE2 in addition to mouse ACE2 addresses SARS-CoV-2’s higher affinity for binding human ACE2, which reduces species tropism and allows virus to cause lethal disease in the mice. As is common in transgenic protein expression, hACE2 under the epithelial cell cytokeratin-18 (K18) promoter is not expressed with the identical tissue localization as it would be in humans. Therefore, viral localization should always be carefully considered. Our observations show that Delta accomplishes lung infection in the mouse and induces strong pneumonia. One popular alternative to the K18-hACE2 mouse model is the Syrian Golden Hamster. After SARS-CoV-2 challenge, hamsters develop more observable respiratory disease and pneumonia due to a higher affinity for hamster ACE2 (64, 65). Delta has been reported to be highly pathogenic in golden hamsters, with increased tropism for the respiratory tract and low development of disease phenotypes like weight loss and drops in temperature (32, 66). The similarities between mouse and hamster models provide evidence for Delta’s unique disease manifestation, defending the importance of its use in preclinical models when compared to prior strains.

With the emergence of the Omicron variant, it’s clear that novel variants of SARS-CoV-2 will continue to arise with enhanced disease capabilities. To continue preclinical COVID-19 research with small animal models, strains that more accurately recapitulate present human disease are necessary. Understanding the unique disease mechanisms of each VOC and the caveats they may present in these models will help to advance research into the identification or improvement of therapeutics.

The K18-hACE2-mouse challenge model of COVID-19 holds high value in characterizing variants, evaluating vaccines and therapeutics, and defining the pathogenic mechanisms of SARS-CoV-2. The data presented here show that a sublethal challenge with Delta induces inflammation that is more histopathological than the Alpha VOC and increases interferon responses. This study also defines the doses of SARS-CoV-2 VOC that are useful for considering in performing preclinical studies. The use of RNAseq allowed us to characterize the entire transcriptome of the mouse lung to understand the effect of challenge by either Alpha or Delta variants. We observed massive differential gene expression caused by Delta despite that fact that the mice did not reach morbidity. Our study identified unexpected and uncharacteristic aspects of Delta’s disease pathogenesis. Together, our study outcomes underscore the need to continue to understand SARS-CoV-2 especially outside of the immunogenicity differences of the spike protein that are most often focused upon. We plan to build upon the findings of this study and characterize single-cell gene expression of the host response in a continued effort seeking to understand ways to combat this virus.

## Materials and methods

### Biosafety, Animal, and Ethics statement

All research performed was approved by West Virginia University IACUC protocol number 2004034204. K18-hACE2-mice were purchased from Jackson Laboratory (B6.Cg-Tg(K18-ACE2)2Prlmn/J; JAX strain number #034860). All SARS-CoV-2 viral propagation or challenge studies were conducted in the West Virginia University Biosafety Laboratory Level 3 facility under the IBC protocol number 20-04-01. SARS-CoV-2 challenged mouse serum and lung supernatant were inactivated with 1% Triton per volume before exiting high containment. Additional tissues were treated using TRIzol reagent or fixed before additional work in BSL2 conditions.

### SARS-CoV-2 cultivation and K18-hACE2 mouse challenge

SARS-CoV-2 strains were obtained from BEI: USA-WA-1/2020 (WA-1; BEI NR-52281) (GenBank accession number: MN985325), hCoV19/England/204820464/2020 (Alpha; NR-54000)(GISAID: EPI_ISL_683466), and hCoV19/South Africa/KRISP-EC-K005321/2020 (Beta; BEI NR-54008) (GISAID: EPI_ISL_678570). The SARS-CoV-2 Delta variant (B.1.617.2 hCoV-19/USA/WV-WVU-WV118685/2021) was obtained from a patient viral transport medium swab (GISAID Accession ID: EPI_ISL_1742834). SARS-CoV-2 strains were propagated in Vero E6 cells (ATCC-CRL-1586). For variant comparison studies, K18-hACE2 mice were intranasally challenged with a volume of 50 uL (25uL per nare) at: 10^3^ PFU/dose (8 week old male and female received WA-1 or Beta; 12 week old female mice received Alpha; 14 week old female received Delta), or 10^4^ PFU/dose (17 week old female received WA-1; 8 week old male and female received Alpha or Beta; 20 week old female received Delta). Male age-matched mice at 8 weeks old were infected with 10^3^ PFU for timepoint comparison study. Viral doses were prepared from the first passage collections from infected Vero E6 cells. Mice were anesthetized with IP injection of ketamine (Patterson Veterinary 07-803-6637) /xylazine (Patterson Veterinary 07-808-1947).

### Disease score of SARS-CoV-2 challenged mice

Challenged K18-hACE2 mice were evaluated daily through both in-person health assessments in the BSL3 and SwifTAG Systems video monitoring for out to 14 days post challenge. Health assessments of the mice were scored based on the following criteria: weight loss (scale 0-5 (up to 20% weight loss)), appearance (scale 0-2), activity (scale 0-3), eye closure (scale 0-2), and respiration (scale 0-2). All five criteria were scored based off a scaling system where 0 represents no symptoms and the highest number on the scale denotes the most severe phenotype. Additive health scores of the criteria listed above were assigned to each mouse after evaluation. Mice that scored 5 or above on the health assessment required immediate euthanasia. Average cumulative disease scoring was calculated by adding the disease scores of each mouse from the group on each day and reporting the mean. Morbid mice that were euthanized during the study, before day 14, retained their disease score for the remainder of the experiment.

### Euthanasia and tissue collection

Mice were euthanized either when assigned a health score of 5 or above or at the end of the experiment with an IP injection of Euthasol (390mg/kg) (Pentobarbital) followed by secondary measure of euthanasia with cardiac puncture. Blood from cardiac puncture was collected in BD Microtainer gold serum separator tubes (Catalog No: 365967), centrifuged at 15,000 x *g* for 5 minutes and serum collected for downstream analysis. Nasal wash was acquired by pushing 1mL of PBS through the nasal pharynx. 500μL of nasal wash was added to 500μL of TRI reagent for RNA purification and the remainder of the nasal wash was frozen for serological analysis. Lungs were separated into right and left lobes. Right lobe of the lung was homogenized in 1mL of PBS in gentleMACS C tubes (order number: 130-096-334) using the m_lung_02 program on the gentleMACS Dissociator. 300μL of lung homogenate was added to 1000μL of TRI Reagent (Zymo research) for downstream RNA purification and 300 μL of lung homogenate was centrifuged at 15,000 x *g* for 5 minutes and the lung supernatant was collected for downstream analyses. Brain was excised from the skull and homogenized in 1mL PBS in gentleMACS C tubes using the same setting as lung on the gentleMACS Dissociator. 1000μL of TRI Reagent was added to 500μL of brain homogenate for RNA purification.

### qPCR SARS-CoV-2 viral copy number analysis of lung, brain, and nasal wash

RNA purification of the lung, brain and nasal wash was performed using the Direct-zol RNA miniprep kit (Zymo Research R2053) following the manufacturer protocol. SARS-CoV-2 copy numbers were assessed through qPCR using the Applied Biosystems TaqMan RNA to CT One Step Kit (Ref: 4392938). We utilized nucleocapsid primers (F: ATGCTGCAATCGTGCTACAA; R: GACTGCCGCCTCTGCTC); and TaqMan probe (IDT:/56-FAM/TCAAGGAAC/ZEN/AACATTGCCAA/3IABkFQ/) that were synthesized according to Winkler. *et al*, 2020 (43). The following final concentrations were used according to the Applied Biosystems TaqMan RNA to CT One Step Kit manufacturer protocol: TaqMan RT-PCR Mix 2X, Forward and reverse primers 900nM final, TaqMan probe 250nM final, TaqMan RT enzyme mix 40X and RNA template 100ng (excluding Nasal Wash). Purified RNA samples with a concentration less than 100ng/uL were not diluted for use in qPCR reactions. All Nasal Wash RNA samples were used at a set volume of 2 uL due to low RNA quantification via the Qubit 3 fluorometer. Triplicates were prepared for each sample, and samples were loaded into a MicroAmp Fast optical 96 well reaction plate (Applied Biosystems 4306737). Prepared reactions were run on the StepOnePlus Real-Time System machine using the parameters: Reverse transcription for 15 minutes at 48°C, activation of AmpliTaq Gold DNA polymerase for 10 minutes at 95°C, and 50 cycles of denaturing for 15 seconds at 95°C and annealing at 60°C for 1 minute.

### Cytokine analysis

R&D 5-plex mouse magnetic Luminex assay (Ref LXSAMSM) was used to quantify cytokines: IL-1β, CXCL13, TNFα, IL-6, IFN-γ, IL-17, and CXCL10 from lung supernatant. Manufacturer protocols were followed in preparing samples. Mouse cytokine plate was analyzed on the Luminex Magpix and pg/mL were calculated based off standard curves generated for each cytokine in the assay. IFN-α, IFN-β and IFN-γ were additionally quantified using MSD U-PLEX Interferon Combo 1 (ms) assay kit (Catalog No K15320K-1) and manufacturer protocols. MSD assay plates were analyzed using the Meso Scale Discovery Sector 2400.

### Histology and Electron microscopy

Left lobes of lungs were immediately fixed in 10mL of 10% neutral buffered formalin. Fixed lungs were paraffin embedded into 5 μm sections. Sections were stained with hematoxylin and eosin and visualized on the DynamyxTM digital pathology platform. Lungs were scored for chronic and acute inflammation in the lung parenchyma, blood vessels, and airways by a blinded pathologist. Pulmonary inflammation was quantified by measuring the total area of lung tissue involved by inflammation. The predominant inflammatory cell type was noted. To quantify vascular margination of inflammatory cells, five representative arteries were identified within areas involved by inflammation. Total length of endothelium of these vessels was measured and the number of marginating inflammatory cells in the cross section were manually counted. Areas involved by inflammation were further evaluated by electron microscopic examination. The areas of interest were punched from the paraffin embedded tissue block and processed for electron microscopy. Ultrathin sections were cut on a Leica Ultra-Microtome, collected on copper mesh grids, stained using uranyl acetate and lead citrate and viewed using a Jeol 1010 electron microscope (FEI) with attached AMT camera.

### Illumina library preparation, sequencing, and *in silico* bioinformatic analysis

RNA quantity was measured with Qubit 3.0 Fluormeter using the RNA high sensitivity (Life Technologies) and RNA integrity was assessed using an Agilent TapeStation. RNA was DNAased before library preparation. Illumina sequencing libraries were prepared with KAPA RNA HyperPrep Kit with RiboErase (Basel, Switzerland). Resulting libraries passed standard Illumina quality control PCR and were sequenced on an Illumina NovaSeq s4 4000 at Admera Health (South Plainfield, NJ). A total of ∼100 million 150 base pair reads were acquired per sample. Sequencing data will be deposited to the Sequence Read Archive. The reads were trimmed for quality and mapped to the *Mus musculus* reference genome using CLC Genomics Version 21.0.5. Two mice were excluded due to no detectable viral reads (n=5 NC, 4 Alpha Day 2, 5 Alpha Day 6, 5 Delta Day 2, 4 Delta Day 6). An exported gene expression browser table is available upon request. Statistical analysis was performed with the Differential Expression for RNA Seq tool and genes were annotated with the reference mouse gene ontology terms. Quantification of the number of activated or repressed genes unique to each experimental group was performed using Venny 2.1 (67). Genes from each experimental comparison with significant fold changes (Bonferroni ≤ 0.04) were submitted to the WEB-based Gene SeT AnaLysis Toolkit’s Over Representation Analysis (ORA) software compared to the reference set “affy mg u74a” to determine GO terms from gene ontology and biological process databases (FDR ≤ 0.05) (68). GO Term heat maps were generated using Morpheus (69). Raw read data is available at NCBI SRA: SUB10957945 (submission complete, pending processing).

### Statistical analyses

All statistical analyses were performed using GraphPad Prism version 9. Statistical analyses were performed with n ≥ 3 for all K18-ACE2 mice studies. Ordinary one-way ANOVA with Dunnett’s multiple comparisons test or Two-Way ANOVA with Tukey’s multiple comparisons test were used with single pooled variance for data sets following a normal distribution and Kruskal-Wallis with Dunn’s multiple comparisons test for non-parametric distributed datasets. Kaplan-Meier survival curves were utilized, and Log-rank (Mantel-Cox) test were used to test significance of survival between sample groups.

## Acknowledgements

This project was supported by the Vaccine Development Center at the West Virginia University Health Sciences Center. F.H.D. and the VDC are supported by the Research Challenge Grant no. HEPC.dsr.18.6 from the Division of Science and Research, WV Higher Education Policy Commission. MSD QuickPlex SQ120 in the WVU Flow Cytometry & Single Cell Core Facility is supported by the Institutional Development Awards (IDeA) from the National Institute of General Medical Sciences of the National Institutes of Health under grant numbers P30GM121322 (TME CoBRE) and P20GM103434 (INBRE). Figure preparation was supported by GraphPad Prism and BioRender.

We would like to acknowledge Mary Tomago-Chesney for sectioning and performing H&E staining on tissues. We would like to thank the Electron Microscopy, Histopathology and Tissue Bank Core for obtaining Electron Microscopy images of the delta variant in lungs. Lastly, we thank Drs. Laura Gibson and Clay Marsh for supporting this COVID-19 vaccine research.

## Author contributions

Studies were designed by FHD, KSL, JRB, AMH. All authors contributed to the execution of the studies. MTW and IM prepared and provided titered viral stocks of WA-1, Alpha, Beta, and Delta SARS-CoV-2 for challenge. Animal health checks, necropsy, and tissue processing were performed by FHD, TYW, BRP, KSL, JRB, OAM, AMH, and HAC. Viral RNA qPCR was performed by HAC and OAM. Serological analysis was executed by KSL, and NAR. Luminex cytokine assays were completed by BPR. MSD assays were performed by KSL. Lung tissue samples were histologically scored by MSA and JPG. Data was analyzed by KSL, AMH, and FHD. All authors contributed to the writing and revision of this manuscript.

